# Polymorphic dynamics of ribosomal proteins gene expression during somatic cell reprogramming and their differentiation in to specialized cells-types

**DOI:** 10.1101/114868

**Authors:** Prashanth Kumar Guthikonda, Sumitha Prameela Bharathan, Janakiram Rayabaram, Trinadha Rao Sornapudi, Sailu Yellaboina, Shaji Ramachandran Velayudhan, Sreenivasulu Kurukuti

## Abstract

Factor induced pluripotent stem cells (iPSCs) offer great promise in regenerative medicine. However, accumulating evidence suggests that iPSCs are heterogeneous in comparison with embryonic stem cells (ESCs), and that is attributed to various genetic and epigenetic states of donor cells. In the light of the discovery of cell-type specialized ribosomal protein composition, its role as the cells transit through different stages of reprogramming and when iPSCs differentiate into specialized cell-types has not been explored to understand its influence in the reprogramming and differentiation process and outcome. By re-analyzing the publicly available gene expression datasets among ESCs, various sources of iPSCs and somatic cells and by studying the ribosomal protein gene expression during different stages of reprogramming of somatic cells and different passages of established iPSCs we found distinct patterns of their expression across multiple cell-types. We experimentally validated these results on the cells undergoing reprogramming from human dermal fibroblasts. Finally, by comparing publicly available data from iPSCs, iPSCs derived specialized cells and it’s *in vivo* counterparts, we show alterations in ribosomal gene expression during differentiation of specialized cells from iPSCs which may have Implications in the context of ribosomopathies. Our results provide an informatics framework for researchers in efficient generation of iPSCs that are equivalent to ESCs.

## Introduction

Pluripotent stem cells (PSCs) have tremendous applications in developmental studies, disease modelling and regenerative therapy [1]. Induced pluripotent stem cells (iPSCs), generated from adult somatic cells by ectopic expression of reprogramming factors, possess properties similar to embryonic stem cells (ESCs), and have been explored as a potential replacement for ESCs in downstream applications [2]. Initially, iPSCs were thought to be very similar to ESCs but later they were found to be substantially different in their gene expression patterns [3]. Irrespective of the source of their donor cell-type, iPSCs were shown to be less efficient in their differentiation potency to other cell-types but were shown to easily differentiate into their respective donor cell-type, highlighting the influence of donor cell-type specific epigenetic memory in this process [4]. However it was noted that continuous passaging of these cells would attenuate these differences between iPSCs of different sources [5]. Like donor cell-lineage specific factors, incomplete DNA methylation, incomplete repression and reactivation of multiple genes [6], persistent donor cell-type specific gene expression or unique gene expression pattern [7] have been attributed to these observed phenomena. The transcriptional/post-transcriptional regulation of these aberrant/unique epigenetic signatures of iPSCs is poorly understood but errors arising during reprogramming or incomplete reversion to pluripotency could be a cause. Since the potential application of iPSCs in regenerative medicine and disease modelling depends on successful cell-type specific differentiation of iPSC, one needs to investigate mechanisms behind reprogramming and differentiation.

In addition to the above mentioned epigenetic determinants and other regulatory components of transcription [2, 8, 9] and post-transcription [10, 11], the components of translation might also influence restricted differentiation of iPSCs. Multiple studies conducted in cell-types ranging from bacteria to malignant cells indicate the existence of ribosomal subpopulations that differ in their protein complement cause diverse functional translational machinery [12-14]. In this regard the occurrence of cell-type specific ribosome composition particularly during generation of iPSCs has attracted our attention. Researchers have reported that ribosome composition is tissue specific and expression levels of different ribosomal proteins (RPs) are different in different tissues/cell-types [15-17]. Interestingly decrease in concentration of a specific RP was shown to affect a spectrum of translated mRNAs without affecting overall protein synthesis in a given cell [18]. This explanation could account for the fact that mutations in some of the RP genes cause abnormality in particular tissue or cell-type, but doesn’t affect the whole body of an organism [19]. Recent mass spectrometric studies on RPs among different cell-types reported by Slavov *et al*., [20] further support the existence of ribosomes with distinct protein compositions and physiological functions. The recent study [21] reveals a more concrete functional link between heterogeneity in ribosome composition and translational circuitry in mouse ESC. Based on these observations, we hypothesized that heterogeneity in cell-type specific ribosome composition could serve as one of the important determinants that might restrict iPSCs to achieve complete pluripotency.

Here, we first analyzed expression pattern of RP genes during different days of reprogramming of four somatic cells to respective iPSCs and compared them with that of human ESC and report distinct patterns in the RP gene expression. Later, we observed these patterns persist in established iPSC lines at extended passages. Finally, we analyzed expression profiles of iPSCs derived specialized cells and their *in vivo* counterparts. Our analysis identified the unusual polymorphic behaviour of various RP gene expressions during this process. These results highlight the importance of ribosome composition in reprogramming of somatic cells and differentiation of iPSCs to specialized cell-types.

## Material and Methods

### Bioinformatics analysis of expression patterns during somatic cell reprograming

The publicly available microarray gene expression data sets (GSE50206) of human ESCs and four different somatic cell-types viz. human dermal fibroblasts (HDF), human astrocytes (HA), normal human bronchial epithelium (NHBE) and human prostate epithelial cell (prEC) that were subjected to reprogramming were analysed [22]. The 75^th^ percentile normalized expression values were downloaded for analysis of RP gene expression. We divided each dataset that consists of ESCs, donor cell and iPSCs derived from a particular donor cell-type into two parts. One with an expression range of -0.5 to +0.5 (range1) and the rest in another part (range2). We extended this to all other cell-types. Next we designed a practical extraction and report language (PERL) program to identify expression state of a given gene. If the gene is expressed the expression level value will be greater than 0.3 will be in range1 and 0.4 in range2 and if the gene is not expressed or down-regulated the value, which is less than -0.2 in range 1 and if it is less than -0.3 in range 2. The thresholds we selected because at these values, the eight expression patterns that were described in this study could be clearly seen under a heat map. Lesser values were not considered as they may hinder the significance of these results. We set minimum threshold to consider a value as “not expressed”/”very less expressed” and an upper threshold to consider as “overexpressed”. We set the parameters for each of the nine expression patterns and applied for this program with same thresholds for all the cell types. The heatmap representing ribosomal genes’ expression (Fig 2) was drawn using Java treeview software tool [23].

### Derivation of iPSC lines and fluorescence activated cell sorting (FACS) of reprogramming cells

Human adult dermal fibroblast was subjected to reprogramming using STEMCAA lentiviral vector using Bharathan et al., (2017) protocol [24]. On day 12 of reprogramming, a single cell suspension was prepared by treating the cells with TrypLE (Gibco). The cells were stained with labeled antibodies, CD13-PE, SSEA-4-Alexaflour647 and TRA-1-60-BV421 (BD Pharmigen) in KOSR based human iPSC medium for 30 minutes at 4^°^C in dark. The stained cells were washed twice with 1X PBS and sorted using FACS Aria III flow cytometer. Based on the co-expression pattern of three markers CD13, SSEA-4 and TRA-1-60, the cells were sorted into four fractions, CD13^+^ve SSEA-4^-^ve TRA-1-60^-^ve, CD13^+^ve SSEA-4^+^ve TRA-1-60^-^ve, CD13^-^ve SSEA-4^+^ve TRA-1-60^-^ve and CD13^-^ve SSEA-4^+^ve TRA-1-60^+^ve. The sorted cells were centrifuged, cell pellet was re-suspended in Tri-reagent and stored at -80^°^C.

### Derivation and establishment different passages of HDF derived iPSCs

The iPSC lines were derived from HDFs by overexpression of OCT4, SOX2, KLF4 and c-MYC (OSKM) using retroviral factor delivery method [2]. The colonies were isolated based on hESC-like morphology, maintained on SNL feeder layers in hiPSC medium and characterized for pluripotency [24]. The fully characterized and established hiPSC lines were maintained in extended cultures on SNL feeders in hiPSC medium and passaged using collagenase-IV treatment. iPSCs representing passage-5 (P-5), P-27, P-43, P-65, P-71 were collected, centrifuged, cell pellet was re-suspended in Tri-reagent and stored at -80^°^C.

### RNA isolation and quantitative PCR analysis

RNA was extracted from fibroblasts, sorted reprogramming cells and iPSC lines using Trireagent (Sigma-Aldrich). 1 μg of total RNA was used for reverse transcription reaction using Primescript RT reagent kit (Takara) according to the manufacturer’s instructions. Quantitative RT-PCR was set up with SYBR Premix Ex Taq II (Takara Bio) using specific RP gene primers (supplementary Table 1) and analyzed on QuantStudio12K Flex (Life Technologies) real-time PCR systems. The raw data was normalized with *ACTB* gene expression.

### Bioinformatics analysis of RNA-seq data sets from iPSC derived adult cells

RNA-Seq data from iPSCs, iPSCs derived specialized cells and their respective *in vivo* counterparts were downloaded from NCBI GEO with accession numbers (Supplementary Table 2) and the expression values were converted to log2 TPM (transcripts per million) [25]. Then the expression values of RP genes and the house keeping genes *ACTB* and *GAPDH* were taken and plotted on heatmap using R.

## Results

### Dynamic expression of RP gene expression during somatic cell reprogramming

Analysis of publicly available microarray gene expression profiles of pluripotent ESCs, different days of reprogramming donor cells-types of NHEB, HDF, HA and PrEC revealed distinct patterns of RP gene expression (Fig 1a (NHEB cells); 1b (HDF cells); 1c (HA cells) & 1d (prEC cells). The time frame of reprogramming process was divided into early, intermediate and late days to define the transitions in expression profile. During this time frame the RP genes were identified to exhibit eight patterns of expression among these four cell-types (Fig 2). In pattern 1, the gene expression levels of reprogramming cells in early days were similar to donor somatic cells and that in late days were similar to that of ESCs (Fig 2a). This indicated that the genes belonging to this category show donor specific expression in the early days and reprogramming factors could easily bring about the shift in expression profile during reprogramming. Gene with pattern 2 retained donor cell-type specific transcriptional profile in early and late days of reprogramming (Fig 2b). The persistent expression pattern of these genes may contribute to donor cell memory in reprogramming cells. Genes exhibiting pattern 3 showed a higher level expression in intermediate days, followed by attaining expression level similar to donor cell-type in late days (Fig 2c). Genes with pattern 4 showed a higher level expression in intermediate days, followed by attaining expression level similar to ESCs in late days (Fig2 d). Genes with pattern 5 exhibited expression level varying from high level in donor cells and reprogramming cells in early days, intermediate level in ESCs and low level in reprogramming cells in late days (Fig 2e). In Pattern 6, gene expression was observed only in the intermediate days of reprogramming and not in ESCs, donor cells or reprogramming cells in later days (Fig 2f). Genes exhibiting pattern 7 were expressed only in reprogramming cells and are not expressed in donor nor in ESC (Fig 2g). Genes with pattern 8 showed expression at the low level in ESCs, intermediate level in donor cells and high in reprogramming cells in late days (Fig 2h). However, it has to be mentioned that expression of a given ribosome protein is not only dynamic but follow different patterns of expression among different cell-types. Overall, most of the genes belonged to Pattern 1 (Fig 2a) wherein the donor cell memory is retained in immediate stages but gets erased over the time and show ESCs type expression at late stage passages. All the genes belonging to various categories in different cell-types described above are listed (Figure 3a).

**Figure 1.**
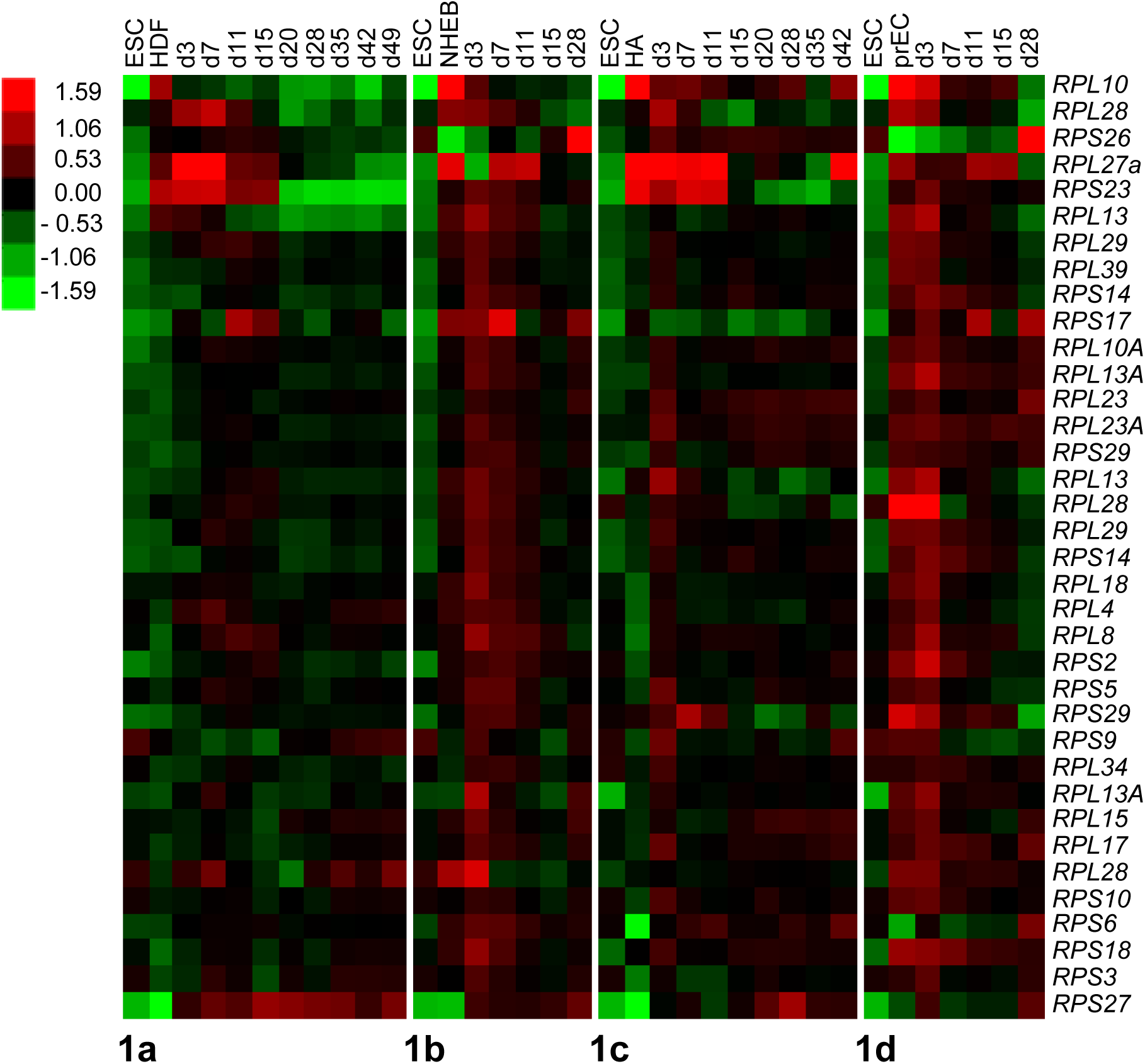
Dynamic expression of RP genes during somatic cell reprogramming: Heat map of group of RP genes showing spectrum of differential gene expression patterns at different days of reprogramming of HDF (**1a**), NHEB cells (**1b**), HAs (**1c**), prEC (**1d**) in comparison with ESCs and respective donor cell-types. Scale bar showing intensity scaled log2 RMA values corresponds to level of expression. Red, black and green colours indicate high, intermediate and low levels of expression respectively. HDF: human dermal fibroblasts (HDF), NHBC: normal human bronchial **e**pithelial cells, HA: human astrocytes, prEC: prostate epithelial cells, ESCs: embryonic stem cells.

**Figure 2:**
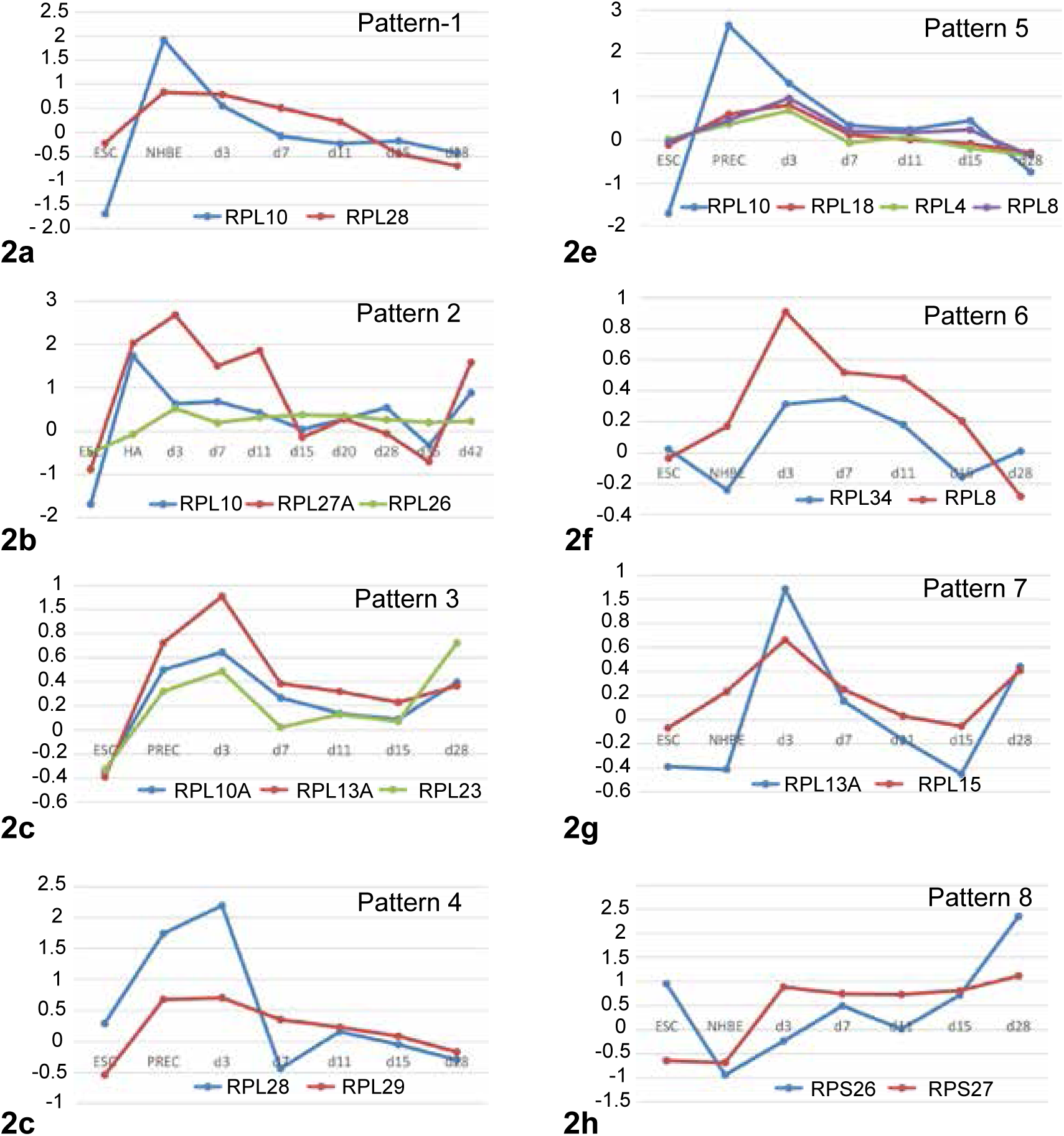
Dynamic patterns (1-8) of RP genes expression during somatic cell reprogramming: Line graphs showing representative RP genes showing pattern 1-8 at different days of either NHEB, HA, prEC somatic cell reprogramming, in comparison with ESCs and respective donor cell-types (2a, b, c, d, e, f, g & h). In patterns-1, expression levels similar to donor cell in early days of reprogramming followed by attaining levels similar to ESCs in late days (**2a**). Pattern-2, expression levels in early and late days, similar to donor cells (**2b**), pattern-3, show higher level expression in intermediate days, followed by attaining expression level similar to donor cell-type in late days (**2c**), pattern-4, show higher level expression in intermediate days, followed by attaining expression level similar to ESCs in later days (**2d**), pattern-5, show higher levels in donor cells and levels decrease in early, intermediate days which are similar to that of ESCs and levels goes further down in later days (**2e**), pattern-6, show higher expression levels only in in the intermediate days of reprogramming but not in ESCs, donor cells or later days of reprogramming cells (**2f**), pattern-7, show expression only in early, intermediate and late days of reprogramming cells but not in donor cell or ESCs (**2g**) and pattern-8, show low levels of expression in ESCs and donor cells but steadily increase in intermediate and late days of reprogramming cells (**2h**) NHBE: normal human bronchial epithelial cells, HA: human astrocytes, PrEC: prostate epithelial cells. ESC: embryonic stem cells. RP genes following representative pattern are depicted in each graph.

**Figure 3:**
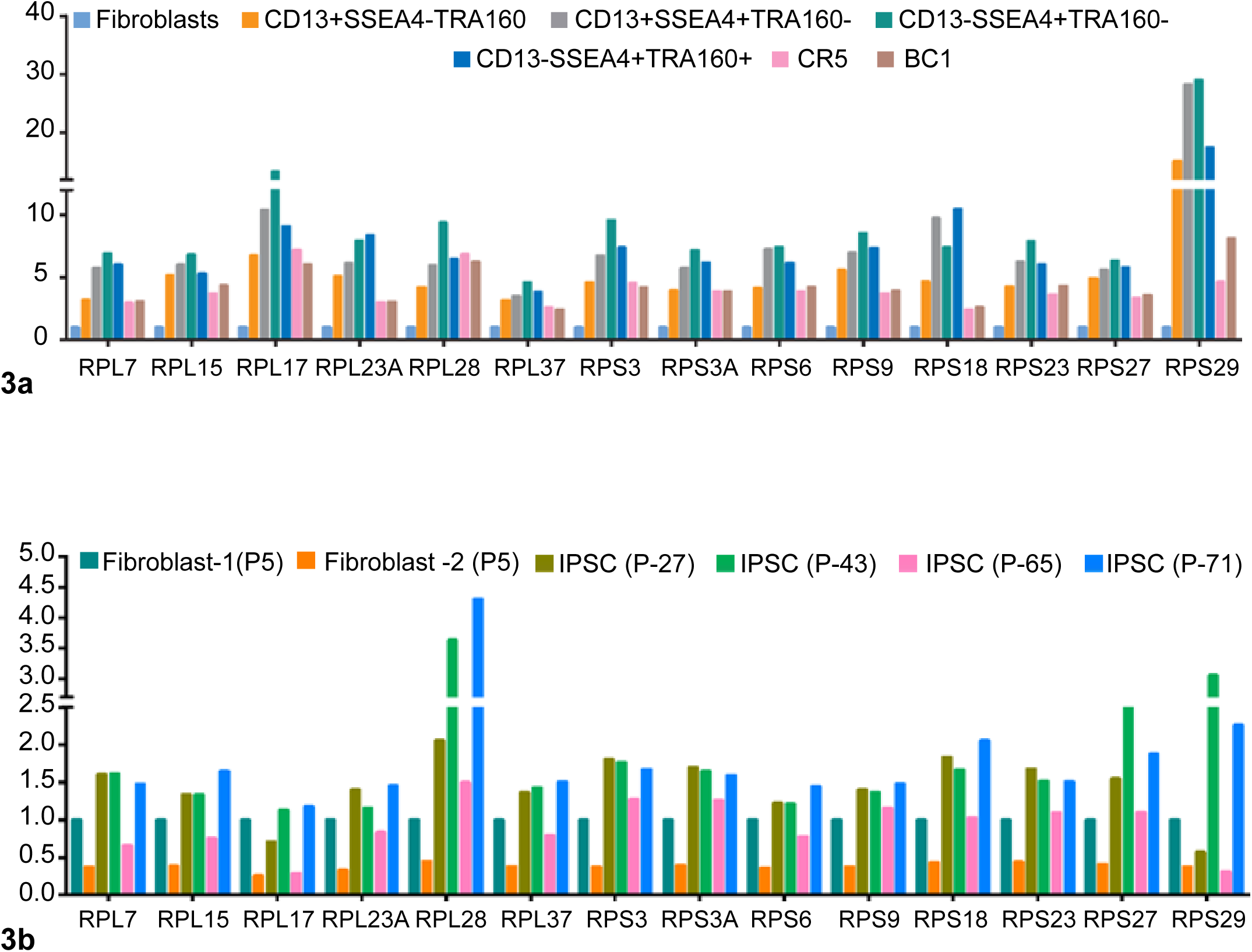
Quantitative PCR validation of RP gene expression during HDF reprogramming and at different passages of established HDF derived iPSCs: (**3a**) Real-time PCR validation of selected RP gene expression during various days reprogramming of HDF in comparison with HDF and established iPSCs (CR5 & BC1) showing peak expression in intermediate stages of reprogramming. (**3b**) Real-time PCR validation of selected RP genes in established hiPSC lines derived from HDFs at different passages (P-5, P-27, P-43, P-65 and P-71), showing dynamic patterns of expression. Values are normalized to *ACTIN-B* and fibroblasts (P-5)-ddCt method (see methods for details).

### Validation of *in silico* observed RP gene expression during somatic cell preprograming

For the validation of *in silico* data on dynamic expression of RP genes, we took the advantage of recently established method of isolation of various stages of OSKM induced reprogramming of HDF by fluorescence activated cell sorting [24] (see methods). The reprogramming cells were sorted based on expression of fibroblast marker, CD13 and pluripotency markers SSEA-4 and TRA-1-60 to obtain cells belonging to different stages of reprogramming namely, fibroblast stage (CD13^+ve^ SSEA4^-ve^ TRA-160^-ve^), intermediate stages (CD13^+ve^ SSEA4^+ve^ TRA-160^-ve^ and CD13^-ve^ SSEA-4^+ve^ TRA-160^-ve^), and late stage (CD13^-ve^ SSEA-4^+ve^ TRA-160^+ve^) (22), and were compared with control iPSC lines, CR5 and BC1 by quantitative PCR. The fourteen RP genes showed varying expression levels in fibroblasts, reprogramming cells at different stages and control iPSC lines (Fig 3a). The fibroblasts showed least level of expression for all analysed RP genes and when subjected to reprogramming, an increase their expression levels were observed, which peaked at the intermediate stage, CD13^-ve^SSEA-4^+ve^TRA-1-60^-ve^. This expression pattern was prominent with *RPL17* and *RPS29*. It was observed that the reprogramming cells at the late stage, CD13^-ve^ SSEA-4^+ve^ TRA-1-60^+ve^ expressed the genes at levels higher than the control iPSC lines. This pattern is clearly evident for *RPL23A*, *RPS9*, *RPS18* and RPS29.

### Dynamic expression of RP genes at various passages of established iPSCs

To check whether these expression patterns continue even after the establishment of induced pluripotency, we analysed established iPSC lines at early (P-27), intermediate (P-43) and late passages (P-65 & 71). The RP gene expression pattern was estimated by quantitative PCR. Interestingly, we observed that most of the genes show similar dynamics during extended culture of iPSC lines and genes like *RPL15*, *RPL17*, *RPL28, RPL37*, *RPS6*, *RPS9* and *RPS18* showed higher expressions at later passages (Fig 3b). Strikingly, these genes show decreased expression from P-43 to P-65 and then gradually show elevated expression in subsequent passage stages P-71 (Fig 3b).

### Dynamic expression of RP gene expression in specialized cells derived from established iPSCs

Differentiation of iPSCs to specialized cell-types is one of the major research focuses in developing iPSCs based regenerative therapy. The RP gene expression patterns in the late passages of the iPSC might influence their differentiation and thereby have an impact on the properties of derived specialized cells. To investigate this possibility, we analysed publicly available RNA seq data for expression patterns of the ribosomal genes in multiple sources of iPSCs derived specialized cells such as neurons and CD34+ hematopoietic stem cells (Supplementary Table 2 & 3). Indeed, we found that the genes such as *RPL7*, *RPL17*, *RPL23A*, *RPS7*, *RPS10* and *RPS27* showed significantly lower expression levels in iPSCs derived specialized neurons when compared with its parental iPSCs and adult neurons (Fig 4a). Other genes such as *RPL28*, *RPL37* and *RPS18* showed similar patterns with lower variations in expression levels [Fig 4a]. These genes were categorized under pattern 8, which were hypothesized to continue their higher expression in the differentiated cells, neurons in this case. More interestingly, genes such as *RPL9*, *RPL10*, *RPL1*4, *RPL24*, *RPL34*, *RPL39*, and *RPS19*, which were not categorized into pattern 8, show a dramatic drop in the gene expression levels compared to iPSC and adult neurons. [Fig 4b] When a similar comparison was made between adult CD34+ hematopoietic stem cells and iPSC (reprogrammed from bone marrow cells) derived CD34+ cells, many genes such as *RPL10*, *RPL13*, *RPL15*, *RPL17*, *RPL21* (Fig4c) showed increased expression in iPSC derived CD4+ cells than their *in vivo* counterpart [Fig 4c] *RPS6KA3,* a protein kinase of *RPS6*, was observed to be deficient in all the somatic cells at late stages of reprogramming. But in the iPSC derived neural cells, there seems to be slightly higher expression than that of neurons.

**Figure 4:**
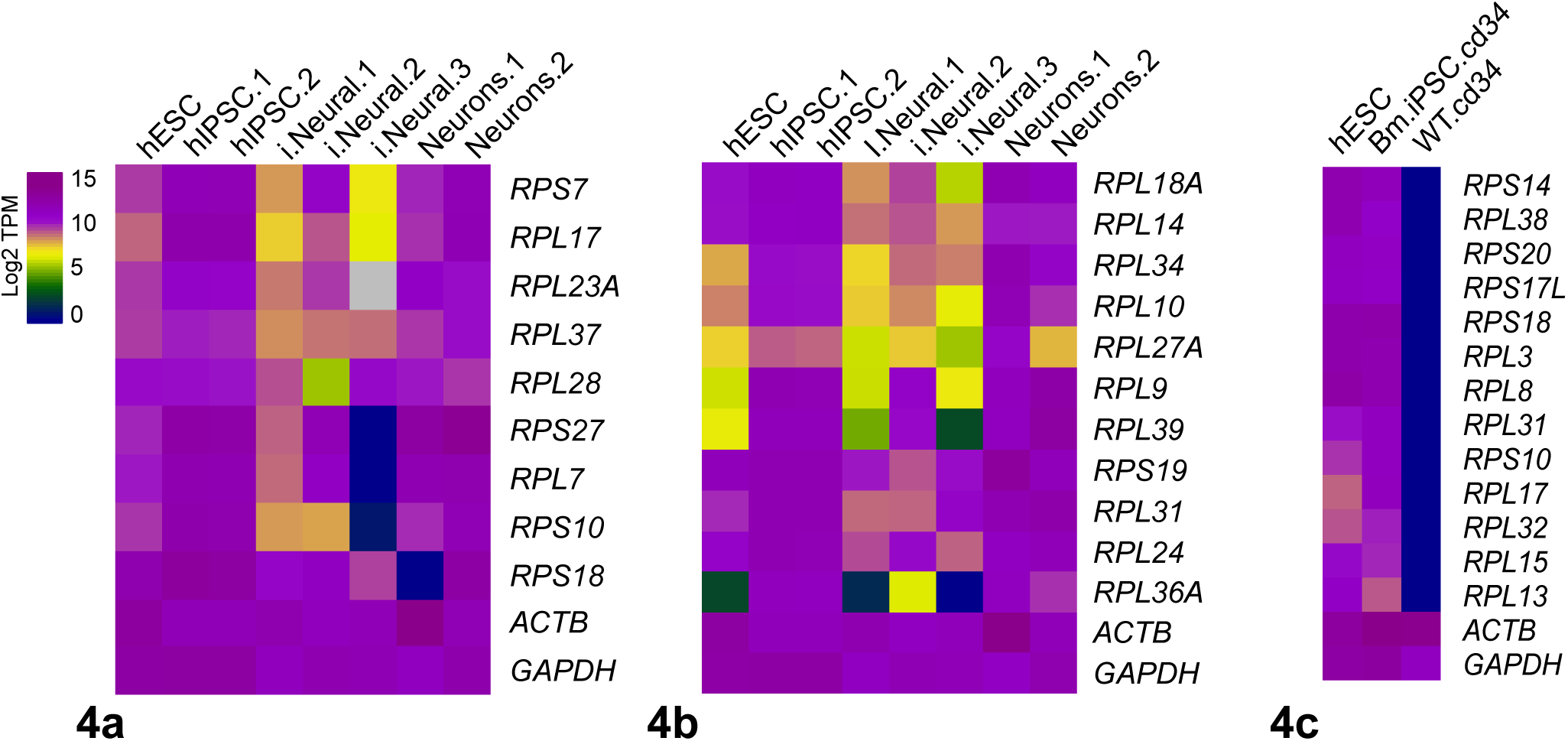
Dynamic expression of RP genes among native and iPSCs derived specialized cells: (**4a**) RP genes which showed pattern-8 of expression during somatic cell reprogramming, showing lower levels of expression in iPSCs derived neural cells than that of its *in vivo* neurons. Expression levels of RP genes in human ESCs and human iPSCs were shown for comparison. (**4b**) RP genes which do not follow pattern-8 of expression during somatic cell reprogramming, showing much lower levels of expression in iPSCs derived neural cells than that of its *in vivo* neurons. Expression levels of RP genes in human ESCs and human iPSCs were shown for comparison. (**5c**) RP genes which show relatively higher levels of expression in iPSCs derived CD4+ve cells than their *in vivo* counterparts. Human ESCs and human iPSCs were shown for comparison.

## Discussion

The observations described in this study in the context of reprogramming of somatic cells and differentiation of iPSCs highlight the importance of RP genes in pluripotency. The RP genes were found to show the distinct pattern of expression during the course of somatic cell reprogramming (Table 1). Hence, regulating the expression of these genes during pluripotency induction may potentially influence the outcome of reprogramming. Abnormal expression of RPs in iPSCs may influence the features of cells differentiated from them and thereby can result in disease phenotypes. The patterns we described in cells during somatic cell reprogramming provide a comprehensive and polymorphic dynamics of RPs gene expression during this process. The RP genes with patterns such as 8 and 9 could be manipulated so that iPSCs will attain expression pattern similar to ESCs. Similarly, further studies on RP genes showing pattern-7 which are expressed highly only in iPSCs, may aid in a better understanding of the process of factor induced reprogramming. Strikingly, these patterns persist in established iPSCs that are maintained in culture for many passages. These patterns give information about the polymorphic behaviour of RP genes dynamic expression in reprogramming cells from four different somatic donor cell-types, which may be helpful in choosing the appropriate donor cell-type for reprogramming and then differentiating them into specialized cell-types.

**Table 1.**
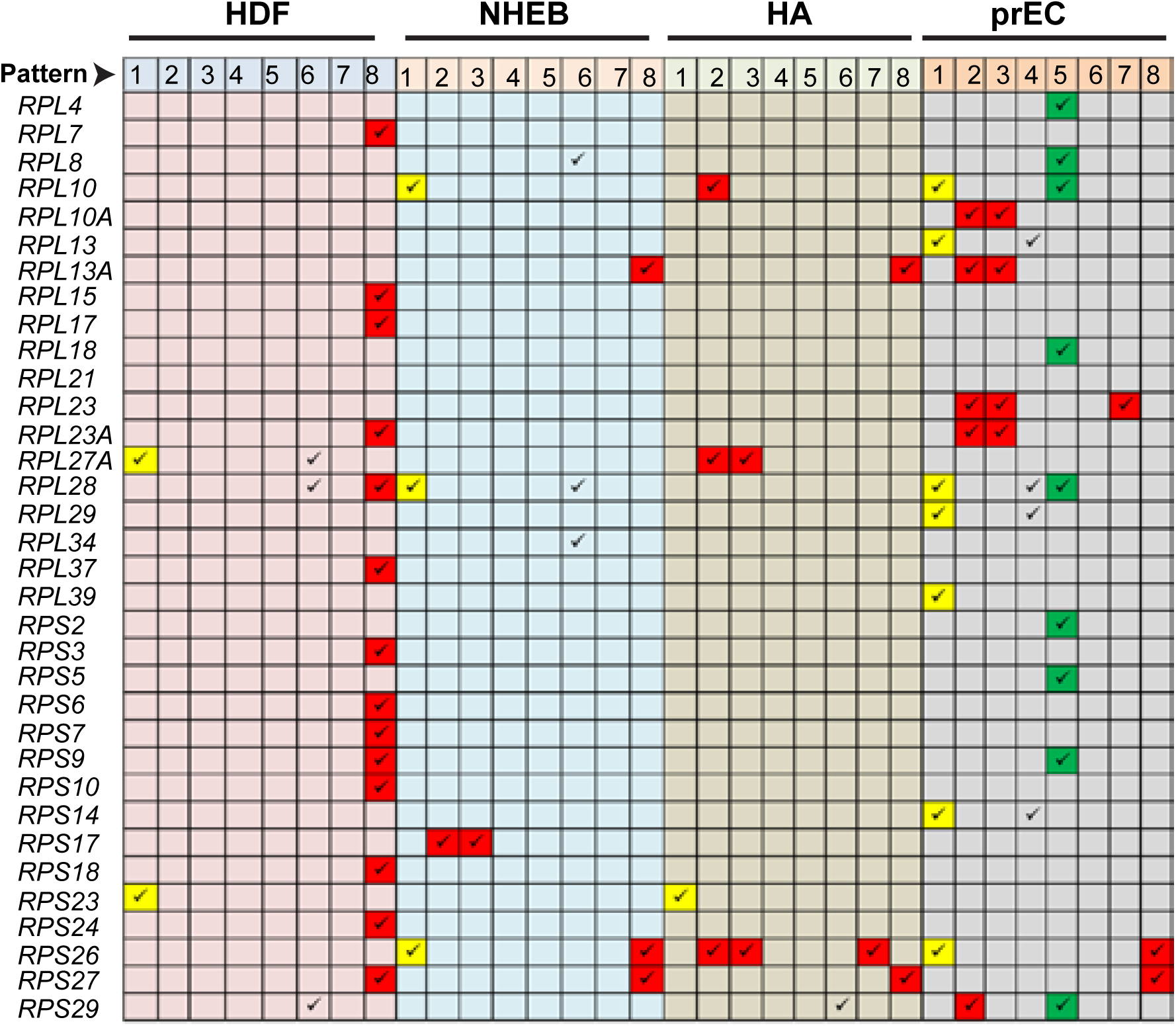
Summary of RP genes following various patterns of expression during somatic cell reprogramming of various donor cell-types: Table showing dynamic pattern of expression followed a specific RP genes either in HDF, NHEB, HA or prEC. Those highlighted in yellow show the typical expected expression pattern upon induction. Those highlighted in red and green were showing variable patterns of expression, which has to be knocked down or over expressed respectively, for efficient reprogramming.

In the protocol which was used to reprogram HDF to iPSC by Takahashi et al [22], they have considered the reprograming up to 49 days from the day of induction and the later days of reprogramming were considered around day-42 and day-49. However, in the protocol by which we generated iPSC from HDF, the later stage of reprogramming is around day-20. This might possibly the reason why our qPCR data during reprogramming is not accurate in accordance with the data from Takahashi et al., [2]. Considering the fact that established iPSCs were reprogrammed only for 20 days in our protocol, these patterns might be again due to the fact that the early passages from established iPSC might be equivalent to the late passages of Takahashi et al., [2]. Despite that, many genes show pattern-8, i.e., elevated expression in later iPSCs, even though the cells were passaged up to 71 times (Fig 3b).

Analysis of specialized cells such as neurons and CD34+ hematopoietic cells derived from iPSCs with their *in vivo* counterparts, the expression patterns of some of the RP genes were found to be different. The patterns in neural cells differentiated from HDF derived iPSCs are different from CD34+ cells differentiated from bone marrow derived iPSCs. The differences we observed here can be partly attributed to the differences in protocols used for factor induced reprogramming and iPSC differentiation or due to inherent cell-type specific genetic and epigenetic differences. In this regard, deficiency of certain RPs in iPSCs derived neurons and CD34+ cells may lead to ribosomopathies. For example, deficiency of *Rpl17* in mouse resulted in enhanced production of shortened 5.8S rRNA [26] Similarly mutations in *Rps7* in mouse is associated with Diamond-Blackfan anaemia (DBA) and neuroanatomical phenotypes [27]and mutations in *RPS19* in humans with DBA [28].

Based on dynamic expression patterns of various RP genes during factor mediated somatic cell reprogramming and at different passages of established iPSCs, it would be predictable that perhaps knocking down selective factors from iPSCs would pave their differentiation towards a specialized cells-types so as to enable them to express similar levels of RP genes as that of its *in vivo* counterparts. However, RP gene expression analysis in specialized cells derived from iPSCs were found to be quite different from both of its *in vivo* counter parts and parental iPSCs themselves, suggesting that heterogeneity in RP genes expression could arise during somatic cell reprogramming, and also during iPSCs differentiation to specialized cell types, reinforcing the fact that one need to carefully evaluate and manipulate their expression profiles before using them for regenerative therapy.

Here in our study, we emphasize the importance of considering the heterogeneity in ribosome composition among various iPSC lines as it can influence their differentiation potential. This study provides a clue that RP composition play an important role in cell-type specific gene regulation and highlights the role of specialized ribosomes in determining the properties of iPSCs. Further elaborate studies need to be conducted to understand the mechanisms of pluripotency and differentiation process of iPSCs for their application in regenerative medicine.

## Conclusions

First, we observed and derived dynamics of Ribosomal proteins’ gene expression during factor induced reprogramming from published datasets. Most of the RP genes in iPSC show similar expression as in that of mESC. Some genes, like in pattern 8, are to be considered for manipulation to obtain expression similar to that of ESC, to avoid the persistent expression in adult cells derived from these iPSC, which may lead to Ribosomopathies. Some of these patterns continued in several passages of iPSC culturing after the establishment of iPSC state. Strikingly, when the expression data from iPSC derived adult cells were observed, many RP genes’ expression is very different from their iPSC as well as from their *in vivo* counterparts. This suggests the need for further studies during generation and differentiation of iPSCs

## Acknowledgments

We would like to thank Ms. Subhalaxmi Mohanthy for her help in PERL programming and Mr. Faizan Ali for his help in analyzing data. Help from BIF facility, School of Life Sciences, University of Hyderabad and FACS facility at Centre for Stem Cell Research, CMC, Vellore are greatly acknowledged. This work was supported by Department of Biotechnology (DBT)-India (BT/PR8688/AGR/36/755/2013) to S.K]. Department of Science Technology-Science and Engineering Research Board (DST-SERB) [SB/YS/LS-230/2013 to S.Y.]; PKG, TRS and SPB acknowledge UGC, CSIR respectively for Senior research fellowship.

## Declarations

Authors declare no conflict of interest

## **Ethics approval and consent to participate**

Not Applicable

## Consent for publication

Not Applicable

## **Availability of data and material**

All data generated or analyzed during this study are included in this published article [and its supplementary information files].

## **Author contributions**

PKG and SK conceived the idea. PKG, TRS & SY analysed the microarray and RNA-seq datasets. SPB, JR & SRV contributed real-time PCR analysis of cells reprogramming from human dermal fibroblasts. PKG, SPB, SRV and SK wrote the paper.

---

**Supplementary Table 1:**
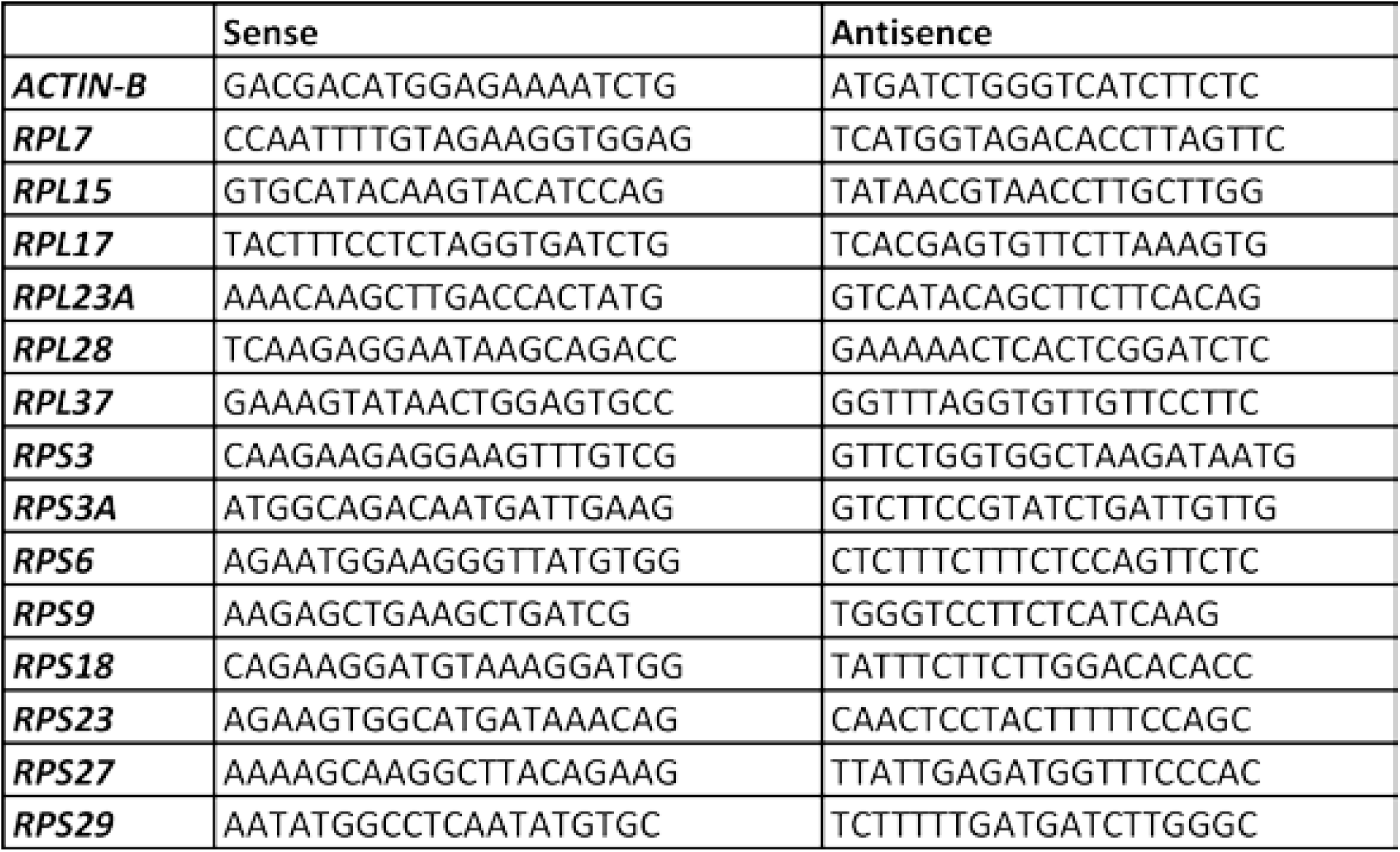
List of RP gene specific primers used in quantitative PCR during different stages of reprogramming HDFs and different passages of established HDF derived iPSCs.

**Supplementary Table 2:** List of publicly available RNA-seq data sets from Gene expression omnibus (GEO) that were used in this study, their origin of cell-types and their respective GEO accession numbers.

| Name | Description | Source |
| --- | --- | --- |
| hESC | Human embryonic stem cells | GSM1536664 |
| hiPSC1 | HDF derived iPSC | GSM2108668 |
| hiPSC2 | HDF derived iPSC | GSM2108669 |
| i-neural_1 | Neuronal cells derived from iPSC of HDF | GSM1536672 |
| i-neural_2 | Neurons derived from iPSC of HDF | GSM2427821 |
| i-neural_3 | Neural cells derived from iPSC of HDF | GSM2507484,GSM2507485,GSM2507486 |
| Neurons1 | Neurons | GSM1585614,GSM1585615 |
| Neurons2 | human peripheral neurons mock sample | GSM2339848 |
| Bm-iPSC-CD34 | CD34+ cells derived from iPSC of bone marrow cells | GSM1464525,GSM1464526 |
| WT-CD34 | CD34+ cells from healthy donor | GSM2754182 |

**Supplementary Table 3:**
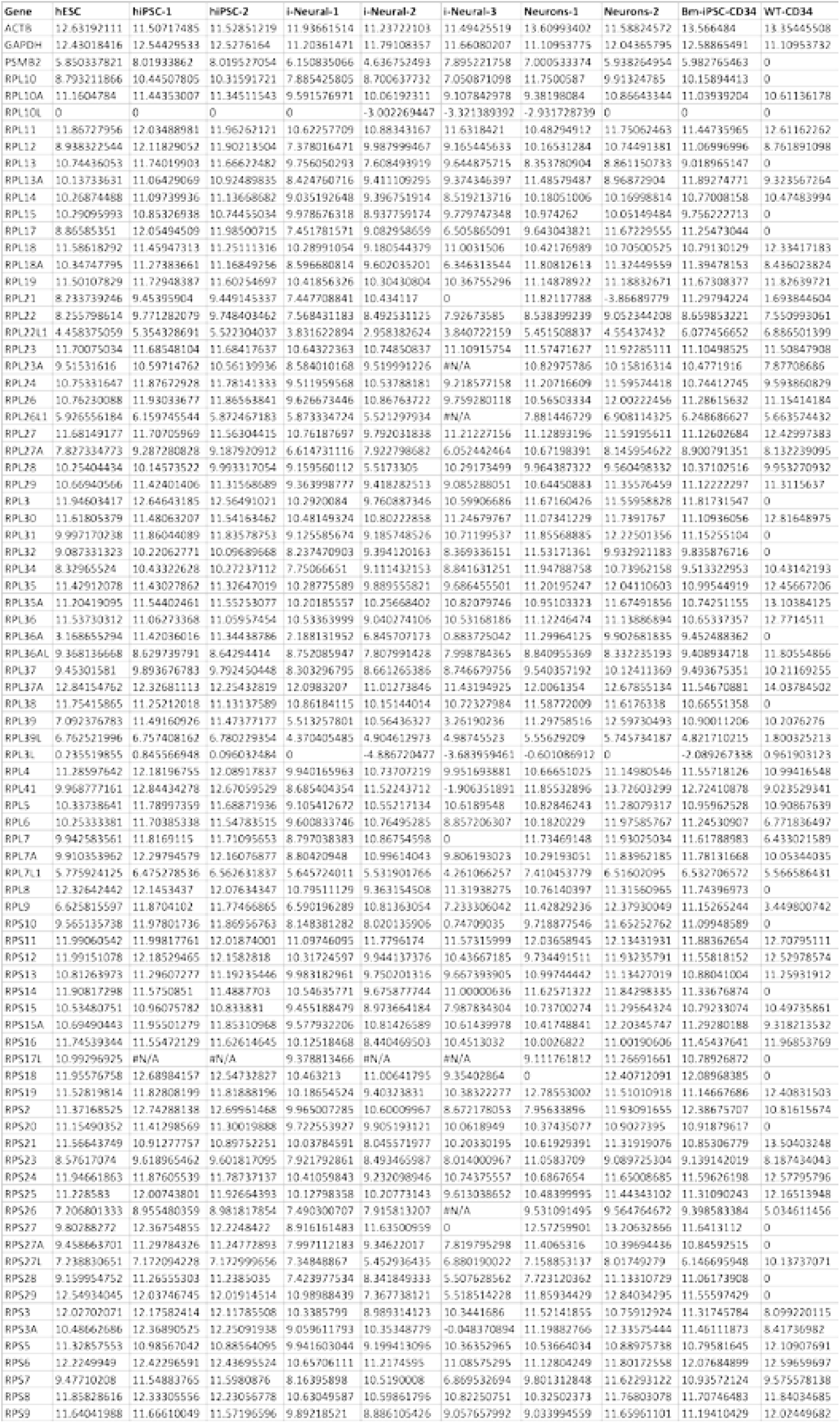
List of all RP genes and their log2 transformed, TPM converted expression values derived from publicly available RNA-seq data sets from hESCs, hiPSCs (1-2), neuronal cells derived from HDF iPSC (i-neural 1-3), i*n vivo* neurons (1-2), CD4 cells derived from bone marrow iPSCs.

## References

1. Keller, G., Embryonic stem cell differentiation: emergence of a new era in biology and medicine. Genes Dev, 2005. 19(10): p. 1129–55.

2. Takahashi, K., et al., Induction of pluripotent stem cells from adult human fibroblasts by defined factors. Cell, 2007. 131(5): p. 861–72.

3. Chin, M.H., et al., Induced pluripotent stem cells and embryonic stem cells are distinguished by gene expression signatures. Cell Stem Cell, 2009. 5(1): p. 111–23.

4. Kim, K., et al., Epigenetic memory in induced pluripotent stem cells. Nature, 2010. 467(7313): p. 285–90.

5. Polo, J.M., et al., Cell type of origin influences the molecular and functional properties of mouse induced pluripotent stem cells. Nat Biotechnol, 2010. 28(8): p. 848–55.

6. Ohi, Y., et al., Incomplete DNA methylation underlies a transcriptional memory of somatic cells in human iPS cells. Nat Cell Biol, 2011. 13(5): p. 541–9.

7. Ghosh, Z., et al., Persistent donor cell gene expression among human induced pluripotent stem cells contributes to differences with human embryonic stem cells. PLoS One, 2010. 5(2): p. e8975.

8. Oldfield, A.J., et al., Histone-fold domain protein NF-Y promotes chromatin accessibility for cell type-specific master transcription factors. Mol Cell, 2014. 55(5): p. 708–22.

9. Cinghu, S., et al., Integrative framework for identification of key cell identity genes uncovers determinants of ES cell identity and homeostasis. Proc Natl Acad Sci U S A, 2014. 111(16): p. E1581–90.

10. Zheng, X., et al., CNOT3-Dependent mRNA Deadenylation Safeguards the Pluripotent State. Stem Cell Reports, 2016. 7(5): p. 897–910.

11. Wang, L., et al., The THO complex regulates pluripotency gene mRNA export and controls embryonic stem cell self-renewal and somatic cell reprogramming. Cell Stem Cell, 2013. 13(6): p. 676–90.

12. Byrgazov, K., O. Vesper, and I. Moll, Ribosome heterogeneity: another level of complexity in bacterial translation regulation. Curr Opin Microbiol, 2013. 16(2): p. 133–9.

13. Deusser, E. and H.G. Wittmann, Ribosomal proteins: variation of the protein composition in Escherichia coli ribosomes as function of growth rate. Nature, 1972. 238(5362): p. 269–70.

14. Guimaraes, J.C. and M. Zavolan, Patterns of ribosomal protein expression specify normal and malignant human cells. Genome Biol, 2016. 17(1): p. 236.

15. Xue, S. and M. Barna, Specialized ribosomes: a new frontier in gene regulation and organismal biology. Nat Rev Mol Cell Biol, 2012. 13(6): p. 355–69.

16. Zhang, W., et al., Decreased expression of ribosomal proteins in human age-related cataract. Invest Ophthalmol Vis Sci, 2002. 43(1): p. 198–204.

17. Bortoluzzi, S., et al., Differential expression of genes coding for ribosomal proteins in different human tissues. Bioinformatics, 2001. 17(12): p. 1152–7.

18. Kondrashov, N., et al., Ribosome-mediated specificity in Hox mRNA translation and vertebrate tissue patterning. Cell, 2011. 145(3): p. 383–397.

19. McCann, K.L. and S.J. Baserga, Genetics. Mysterious ribosomopathies. Science, 2013. 341(6148): p. 849–50.

20. Slavov, N., et al., Differential Stoichiometry among Core Ribosomal Proteins. Cell Rep, 2015. 13(5): p. 865–73.

21. Shi, Z., et al., Heterogeneous Ribosomes Preferentially Translate Distinct Subpools of mRNAs Genome-wide. Mol Cell, 2017. 67(1): p. 71–83 e7.

22. Takahashi, K., et al., Induction of pluripotency in human somatic cells via a transient state resembling primitive streak-like mesendoderm. Nat Commun, 2014. 5: p. 3678.

23. Saldanha, A.J., Java Treeview--extensible visualization of microarray data. Bioinformatics, 2004. 20(17): p. 3246–8.

24. Bharathan, S.P., et al., Systematic evaluation of markers used for the identification of human induced pluripotent stem cells. Biol Open, 2017. 6(1): p. 100–108.

25. Wagner, G.P., K. Kin, and V.J. Lynch, Measurement of mRNA abundance using RNA-seq data: RPKM measure is inconsistent among samples. Theory Biosci, 2012. 131(4): p. 281–5.

26. Wang, M., et al., Reduced expression of the mouse ribosomal protein Rpl17 alters the diversity of mature ribosomes by enhancing production of shortened 5.8S rRNA. RNA, 2015. 21(7): p. 1240–8.

27. Watkins-Chow, D.E., et al., Mutation of the diamond-blackfan anemia gene Rps7 in mouse results in morphological and neuroanatomical phenotypes. PLoS Genet, 2013. 9(1): p. e1003094.

28. Koga, Y., et al., Reduced gene expression of clustered ribosomal proteins in Diamond-Blackfan anemia patients without RPS19 gene mutations. J Pediatr Hematol Oncol, 2006. 28(6): p. 355–61.

